# Repetition-attenuation of auditory N1 is modulated by the phonological representations of spoken word-forms

**DOI:** 10.1101/2021.10.13.464244

**Authors:** Jinxing Yue, Peng Wang, Jiayin Li, Zhipeng Li, Xia Liang, Yifei He

## Abstract

Repeated auditory stimuli are usually found to elicit attenuated peak amplitude of the N1 component of the event related brain potential (ERP). While the repetition-attenuation of the auditory N1 has been found sensitive to some cognitive factors, less is known whether and how the representational properties of stimuli influence this physiological phenomenon. To further address this issue, we focus on the phonological representations of spoken word-forms, and hypothesise modulatory roles of two phonological features: the lexicality and its usage frequency of a word-form. To test this, we used a short-term habituation design with a factorial combination of the two features at two levels each (i.e., lexicality (real versus pseudo word-form) × frequency (high versus low frequency)). EEG was recorded from 30 native Mandarin-speaking participants while they were passively delivered with stimulations trains. Each train consisted of five presentation positions (S1 ∼ S5), on which one word-form is presented repeatedly, separated by a brief, constant interstimulus interval. At the fourth presentation position (S4), we found greater N1 attenuation in low-frequency pseudo word-forms than in low-frequency real and high-frequency pseudo word-forms, respectively. The results support our representational modulation hypothesis, and provides the first evidence that representations of different phonological features interactively modulate the N1 repetition-attenuation. The brain function that underlies the phonological effects of the representational modulation on N1 repetition-attenuation might be sensory filtering.

## 1. Introduction

Neural responses to repeated stimulations at brief intervals tend to decrease. Such adaptive neural mechanism has been termed as habituation, repetition suppression, sensory gating, or, more generally, neural adaptation (e.g., Boutros et al., 2011; Budd et al., 2013; Matsuzaki et al., 2012; Rosburg et al., 2004). It is considered as a basic manifestation of neural plasticity, or a form of learning which forms the foundation for developing more complicated neural plasticity (Garrido et al., 2009; Groves & Thompson, 1970; Mildner, 2008).

In the auditory domain, repetition-induced attenuation of neural responses is already observable at the earliest latencies, as indexed by auditory N1 component of the event related brain potential (ERP) (Fruhstorfer et al., 1970; Woods & Elmasian, 1986), typically recorded around the vertex (Fruhstorfer, 1971; Fruhstorfer et al., 1970; Muenssinger et al., 2013; Rosburg et al., 2002, 2010), by using electroencephalography (EEG) and magnetoencephalography (MEG) for its magnetic equivalent N1m (or N100m) (Näätänen & Picton, 1987; Teismann et al., 2004). The EEG and MEG are known to offer high temporal resolution at an accuracy level of millisecond, and thus, are ideal tools to capture the neural activities related to early sensory processing. Auditory N1 (or N100) ^1^, an evoked response component, is identified as the first prominent negative deflection, peaking around 100 ms after a stimulus onset. N1 and its preceding and following positive components P1 and P2 consist of the P1-N1-P2 complex of ERPs, reflecting a summed synchronous firing of auditory neurons, indicating obligatory, pre-attentive auditory processing even outside the focus of attention (Joos et al., 2014).

The N1 repetition-attenuation has been classically considered as the result of neural refractoriness which reflects *pure* bottom-up adaptation given repeated stimuli (Budd et al., 1998; Ritter et al., 1968; Rosburg et al., 2010). A possible physiological account for this process is that during repeated stimulation, the neurons are subjected to depletion of recyclable neurotransmitters to generate response with the same magnitude as that to the first stimulation (Sara et al., 2002; Wang & Knösche, 2013). According to this refractory account, the N1 repetition-attenuation between acoustically similar stimuli should be alike, because of the highly overlapping neural representations of these stimuli (Marklund et al., 2020).

Besides this *pure* refractory account, there are also findings suggesting some top-down mechanisms in modulating the repetition-attenuation of auditory evoked responses (Font-Alaminos et al., 2020; Herrmann et al., 2018; Öhman & Lader, 1972; Todorovic et al., 2011). For example, Todorovic et al. (2011) found that compared to expected stimulus repetition, unexpected repetition of a pure tone led to stronger neurophysiological responses from 100 ms post stimulus onset. This result suggests an expectation-driven modulation on the attenuation of neural response in the N1 time window, during early auditory perception.

Furthermore, the repetition-attenuation of N1 has been found to differ between speech and non-speech stimuli (e.g., Marklund et al., 2020; Teismann et al., 2004; Woods & Elmasian, 1986). For example, Woods and Elmasian (1986) found more attenuated N1 responses to repeated speech sounds (synthesised vowels) than to non-speech sounds, which are matched with major psychophysical features, such as loudness, duration, and formant frequencies. In contrast, two more recent studies reported less attenuated N1 to repeated speech sounds than non-speech sounds, with careful controls for the acoustic features (Teismann et al., 2004; Marklund et al., 2020). Despite the opposite attenuation directions in speech and non-speech sounds, these studies seem to suggest that the repetition-attenuation of N1 is modulated according to stimuli’s representational properties, instead of their basic acoustic parameters.

As an interim conclusion, when the same auditory stimulus is repetitively presented with a brief interval, attenuated N1 amplitudes are not only the result of a *pure* neural refractoriness but also modulated by a top-down mechanism which is related to the cognitive and/or representational properties of auditory stimuli (Figure 1).

**Figure 1.**
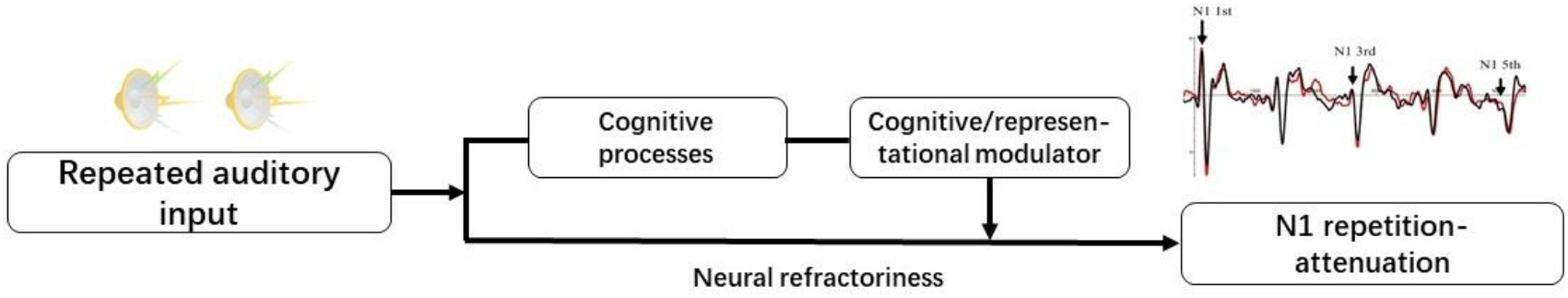
Demonstration of the cognitive/representational modulation on N1 repetition-attenuation induced by repeated auditory stimuli. When cognitive/representational factors are not involved in the perception of repeated auditory stimuli, N1 repetition-attenuation may be the result of neural refractoriness. However, the processing of cognitive/representational features of the input can activate the modulator which influences the outcome of N1 repetition-attenuation.

The representational-modulation conclusion has been tested in a recent study by using stimuli of spoken word-forms (Yue et al., 2017). In this study, researchers hypothesised differential N1-attenuation patterns between the real-word and the pseudo-word condition, given that their representations are at different levels. Real words are meaningful lexical units, and thus, occupy lexical-level representations of phonology. In contrast, pseudowords are word-like speech forms which obey the phonological rules of a language, but do not have corresponding semantic representations (e.g., *bite* versus pseudoword \**bipe*^2^ in English, *cf*. Gansonre et al., 2018; Shtyrov et al., 2017). Hence, pseudowords are believed to be processed at a sublexical level. The representational distinction between real words and pseudowords have made them ideal linguistic models to distinguish cognitive and neural activities that are specific for the lexical-level representations from those to the lower, sublexical-level representations (e.g., Howard et al., 1992; Myers & Blumstein, 2008; Orfanidou et al., 2006; Pulvermüller et al., 2001).

Using two monosyllabic Mandarin spoken word-forms, the researchers (Yue et al., 2017) compared the degree of N1 repetition-attenuation throughout four-time repetitions of a real word (/ma1/) and that in repeated presentations of an acoustically similar pseudoword (*/na1/). They observed greater N1 repetition-attenuation ratio in the pseudoword condition than in the real-word condition, in a region of interest over the right-hemispheric, frontocentral scalp, with possible confounding effects of the phonetic contrast between the real and pseudo word-forms being controlled. Given that the real word differs from the pseudoword in owning the lexical-level representation, this N1 repetition-attenuation effect of lexicality is direct evidence that the repetition-attenuation of N1 is sensitive to the hierarchical difference between phonological representational levels.

Based on these previous studies, here, we aim at further testing the degree to which the repetition-attenuation of N1 is modulated by the representational properties of an auditory word. Besides lexicality (real word vs. pseudoword) as in Yue et al. (2017), we also test if the usage frequency of a word-form modulates the N1 repetition-attenuation. To date, few studies have examined how N1 repetition-attenuation responds to multiple representational properties of phonology. However, the neurophysiological literature has repeatedly suggested that phonological features of spoken words are indeed processed in pre-attentive speech perception. For example, by measuring the Mismatch Negativity (MMN), researchers have found evidence of pre-attentive processing of phonological features such as lexicality and lexical frequency. The lexicality effect of MMN is usually shown as a higher amplitude of the MMN to real words than to acoustically matched pseudowords (e.g., Pulvermüller, et al. 2001, Jacobsen et al., 2004); the lexical-frequency effect suggests greater MMN responses to high-frequency words than to low-frequency words (Aleksandrov et al. 2020; Shtyrov et al., 2011). Like auditory N1, MMN is an obligatory, sensory response elicited by stimuli with a lower probability of recurrence (deviants) in an auditory stimulation sequence, which contains another stimulus with a much higher recurrence probability (standards) (Näätänen, 1995). It presents as a negative deflection, peaking between 100 – 250 ms post stimulus onset, which is partly overlapped with the typical time window for the N1 wave. Moreover, the phonology-tuned MMN responses are considered to indicate highly automatic phonological processing since they can be elicited outside the focus of participant’s attention (Shtyrov & Pulvermüller, 2007).

Moreover, a few studies also found that lexicality and frequency may even interact with each other during pre-attentive speech perception (Shtyrov et al., 2011; Silva et al., 2019). For example, Silva et al. (2019) found that only pseudowords which were generated based on low frequency words elicited less negative N1-P2 responses than the real-word baseline, but pseudowords based on high-frequency words did not.

Hence, according to the cognitive/representational modulation conclusion (See Figure 1), if the phonological features can be processed automatically in pre-attentive speech perception, it can be hypothesised that the N1 repetition-attenuation in spoken words is subjected to modulations of representations of these features (the lexicality and the usage frequency of word-forms). To test this, we adopt a short-term habituation design with monosyllabic spoken word-forms in Mandarin Chinese to elicit attenuated N1 responses.

Monosyllabic words in Chinese are basic lexical units which can either function as words or morphemes to build up polysyllabic words (e.g., [huo4](豁) means wide, [da2](达) means access, and [huo4da2](豁达) means being open-minded, in Gu et al., 2012). It is usually comprised with an onset (initial consonant), a rime (a vowel or a nasalised vowel with /n/ or /?/) and a tone^3^ (Duanmu, 2002). Taking advantage of the special phonological system of Mandarin, the lexicality of a word-form is manipulated by combining the same segmental template with different tones, yielding an existent lexical unit (e.g., /tun1/ (吞), /tun/+Tone1, means to swallow), or a meaningless pseudo word-form (e.g., */tun3/, /tun/+Tone3) (see Yue 2016 for the investigations of the distinctions between the two types of word-forms with behavioural and electrophysiological measures). The same segmental parts between the real and pseudo word-forms ensure that the potential lexicality effect does not involve any processing of anomalous segments (e.g., rime or onset), which cannot be avoided by using pseudowords in non-tone languages, such as English and French, derived through manipulating phonemes in a real word (e.g., Dufour, 2008; Slowiaczek & Hamburger, 1992).

Meanwhile, the usage frequency of meaningful, real word-forms is manipulated, which is measured by the phonological frequency, computed by adding the total frequency of all possible words or morphemes sharing the same segment-tone pattern (*cf*. Ziegler et al., 2000). A monosyllabic Chinese word-form is known to have a number of homophones, which share the same segment-tone pattern (e.g., /yi4/ has homophones like 义, 意, and 亿). Therefore, for a spoken word-form presented in an isolated way, the phonological frequency of a word-form is more practical in reflecting the representations corresponding to the probability of such sound pattern occurs in a language than lexical frequency of a word. Furthermore, the pseudo word-forms inherit the phonological frequencies from the real words whose tones are changed to derive the pseudowords (*cf*. Balota et al., 2004; Perea et al., 2005), because the experience with the use of real words determines how the segmental templates are represented as sublexical units.

The effects of lexicality and frequency are examined to test their hypothesised modulations on N1 repetition-attenuation. If the representations of the two phonological features are independent modulators, only should main effects of lexicality and/or frequency be observed. If they interplay with each other in modulating N1 repetition-attenuation, interaction between the two factors should be identified. Otherwise, if the two factors play no roles in influencing spoken-word elicited N1 repetition-attenuation, no effects of the two phonological features should be found, indicating that N1 repetition-attenuation is a *pure* neural refractory process.

## 2. Method

### 2.1. Participants

Thirty right-handed (adapted Edinburgh Handedness Inventory, Oldfield, 1971) native Mandarin speaking participants (Age mean = 21.7, SD = 4.4; 16 females) who reported no hearing or language disorders were paid to participate in the study. To ensure that they have homogenous language background, they were all born and grown up in the north-eastern region of China, where people are known to speak Mandarin Chinese. They were randomly assigned to one of the two lists of test materials. The experiment was approved by the Ethical Committee of School of Management of Harbin Institute of Technology. Informed consent was given before the experiment according to the Declaration of Helsinki.

### 2.2. Design and Materials

Spoken word-forms in four conditions were generated by combining usage frequency with two levels (high or low), and lexicality of the monosyllabic word-forms stimuli with two levels (real- or pseudo-word). The four conditions are 1) high-frequency real word-forms (HPRW), 2) low-frequency real word-form (LPRW), 3) high-frequency pseudo word-form (HFPW), and 4) low-frequency pseudo word-form.

Forty word-forms were derived from twenty segmental templates by combining one of which with two different tones, leading to a real and a pseudo word-form, respectively (e.g., /gei3/ ‘给’ (to give) and */gei1/). The frequency data were from the Chinese Internet Word Frequency List of the Lancaster Corpus of Mandarin Chinese (McEnery & Xiao, 2004). Segmental templates that appear more than 100 times per million words in the database were considered to be of high frequency, otherwise low frequency (*cf*. Shu et al., 2003; Zhang et al., 2009).

Considering that tone regularity is a factor influencing spoken word recognition (Wiener & Ito, 2015; Wiener & Turnbull, 2016), all real word-forms included in this experiment were with high regularity, that is, the most regularly combined tone given a segment template. For instance, [mai3] (e.g., to sell) was chosen because it is more regular than other lexical combinations, such as [mai2] (e.g., to burry), even though they are all meaningful word-forms. This also makes sure that the phonological frequency of a real word-form is representative enough for its tone-manipulated pseudo counterpart. To avoid presenting the same segments in more than one condition to the same participant, materials were distributed into two lists following the Latin Square method. In addition, each list also contained 10 real and 10 pseudo word-forms as foils. The stimuli were previous recorded for a series of lexical decision experiments by using high quality recording equipment, articulated by a female, native Mandarin speaker (Yue, 2016). Stimuli were normalised for the same average intensity (75 dB) and duration (450 ms) by using an acoustic software programme, PRAAT (Boersma & Weenink, 2013).

A short-term habituation paradigm was employed to induce repetition-attenuation. A stimulus was programmed to be presented in stimulation trains, each of which held five repeated presentation positions (S1 to S5), separated with a constant inter-stimulus interval (ISI) for 450 ms. Given an inter-train interval (ITI) for 5s with a 200-ms jitter, the electrophysiological response to a spoken word-form could be expected to recover from the attenuation of N1 in the previous train, known as a critical characteristic of short-term habituation (Rosburg et al., 2002, 2010). The word-forms in each list were divided into five blocks, in which one word-form out of five in each condition only appeared in one block. In a block, a train carrying the same word-form was delivered 11 times. Besides, four foil word-forms were included and varied for each block but kept the same for the two lists. The train of a foil word-form was presented five times in a block. The trains of stimuli were pseudo-randomly presented for those of the same condition were presented no more than three times in a row. A 1.5 min stimulus-free break was set between every two blocks.

### 2.3. EEG data acquisition

The EEG of each participant was recorded in a sound attenuated cabin with constant lightness, seated in front of a PC monitor placed at a distance of about 1.2 m. They were randomly administered to be exposed to one list of stimuli, and were suggested to refrain from unnecessary body movement. During the EEG recording, auditory stimuli were delivered binaurally and passively via a pair of Sennheiser headphones. To avoid potential confounds of selective attention on the auditory habituation of evoked potentials (Öhman & Lader, 1972), participants were instructed to watch a silent cartoon movie without subtitles, and asked to remember the contents for a good performance in a comprehension test which was administered after the EEG experiment. The comprehension test required participants to read 12 statements about the movie, and judge whether they matched the contents of the movie by choosing either “Yes”, “No”, or “Cannot remember”. Moreover, they were encouraged to ignore any sounds that would be presented from the headphones.

The EEG signal was recorded by a LiveAmp amplifier (Brain Products) via 32 Ag/AgCl electrodes situated on an elastic cap, according to the extended international 10-20 system, with a 500 Hz sampling rate. One electrode was placed at the right infraorbital ridge to monitor ocular movement. The online reference was FCz and the ground electrode was AFz. The impedance of electrodes was kept below 5 kΩ.

### 2.4. EEG data analysis

EEG pre-processing was performed in Brain Vision Analyzer 2.0 (Brain Products). The EEG data were band-pass filtered between 1 and 30 Hz, and re-referenced to the average amplitudes recorded from the two mastoids. Voltage moves higher than 40μv were recognized as artefacts and rejected for further data analysis. Then, the EEG recording was segmented separately for the five presentations (i.e., S1∼S5) of each stimulus type, with an epoch of 600 ms, between −100 ms before and 500 ms after the stimulus onset. Baseline was corrected according to the pre-stimulation responses.

Peak amplitudes of N1 were quantified with a peak-to-peak measurement, by calculating the amplitude difference between an N1 peak and the peak amplitude of the first prominent positive component P1 that precedes the N1 (*cf*. McClaskey et al., 2018; Toyokura, 2003). For this measurement, peak amplitudes of P1 and N1 were detected in two time-windows, respectively: 40-100 ms (P1) and 80-170 ms (N1) for each type of stimuli at every presentation position, per participant. The time windows were defined by referring to previous studies (e.g., Swink & Stuart, 2012; Grau et al., 2007) and adjusted according to visual inspection of the data. The degree of N1 repetition-attenuation at each presentation position was quantified by attenuation index, which is calculated as the ratio of the N1 peak amplitude in a repeated position (S2 ∼ S5) to that in the initial presentation (S1) or each of the four stimulus types (i.e., Attenuation index = N1_Sn_/N1_S1_, in which n = 1, 2, 3, 4, 5). Accordingly, the greater the index is, the less the relative neural attenuation takes place.

The N1 repetition-attenuation was analysed at two representative electrodes, namely C3 and C4. The selection of the two electrodes were based on three criteria, defined *a priori*. First, they should be around the vertex electrode Cz to reflect typical N1 repetition-attenuation (Rosburg et al., 2010). Second, electrodes from the right fronto-central scalp region should be included, in order to replicate the lexicality effect in Yue et al. (2017). Third, electrodes from the left fronto-central area should be chosen as they usually capture pre-attentive neural processing of speech (Pulvermüller & Shtyrov, 2006). Putting these together, we selected C3 and C4 situated in the left- and right-hemispheric scalp regions to investigate the hypothesised modulation of speech phonology on N1 repetition-attenuation. Since the two electrodes are far from eyes, the signals are less contaminated by ocular movements, and thus, no ocular correction was performed except artifact rejection. Only less than 4% of trials at the two electrodes were rejected as artefact given the strict rejection criteria.

The data analysis began by testing the validity of the current paradigm in eliciting reliable N1 repetition-attenuation. A 5×2×2×2 repeated measures ANOVA with four factors: PRESENTATION (S1 ∼ S5), LEXICALITY (Real vs pseudo word-forms), FREQUENCY (high versus low frequency), and ELECTRODE (C3, C4) was first conducted to examine whether the N1 at S1 differs from the other repeated presentations, which could be verified by a main effect of PRSENTATION.

Inferential statistical analyses were conducted with the attenuation-index data at the four repetition positions, separately, beginning with a 2×2 ×2 repeated measures ANOVA with three factors: LEXICALITY, FREQUENCY, and ELECTRODE. Repetition position was not considered as a factor because the degree of attenuation may not be a linear function of presentation positions (Rankin et al., 2009), and thus, the presentation position (i.e., PRESENTATION) may not be a suitable factor for ANOVA. If any interactions between LEXICALITY and FREQUENCY could be identified, further analyses would be carried out to check how the factors lead to differential attenuation patterns in the four types of stimuli. Greenhouse-Geisser correction was applied to adjust the degrees of freedom for the *F* tests when the sphericity assumption was not attested by the Mauchly’s Test. Bonferroni corrections were performed when appropriate. The uncorrected degrees of freedom and the adjusted *p*-values were reported. The significance criterion was *p* < .05.

## 3. Results

N1 repetition-attenuation was successfully elicited with the current paradigm as reflected by the visual inspection of apparent decrement of the N1 responses in repeated stimuli (S2 – S5) relative to S1 (see Table 1, also see Figure 2 and 3 for demonstration of the attenuated amplitudes of the early ERP components in S4 relative to those in S1 and the N1 topography in S1 and S4). This response pattern is confirmed by a significant main effect of PRESENTATION (*F* (4, 116) = 32.55, *p* < 0.001, partial *η*^2^ = 0.529). This result attests the validity of the short-term habituation design used in this study. Moreover, the participants had very high accuracy rates in the statement judgement task (M = 90.4%, SD = 7.1%), suggesting that they have focused on the movie-watching task, and therefore, can be assumed to spare little attention on the auditory stimuli. A summarisation of the N1 peak amplitudes and the attenuation indices can be seen in Table 1.

**Table 1.** Peak-to-peak amplitudes of the N1 amplitude (i.e., measured as P1-N1 peak-to-peak amplitude) at S1, and the N1 repetition-attenuation ratios for the four repeated positions in the two electrodes of interest (C3 and C4)

| Condition | Electrode | Amplitude S1 ($\mu\text{v}$) | | Attenuation Index S2 | | Attenuation Index S3 | | Attenuation Index S4 | | Attenuation Index S5 | |
| --- | --- | --- | --- | --- | --- | --- | --- | --- | --- | --- | --- |
|  |  | *M* | *SD* | *M* | *SD* | *M* | *SD* | *M* | *SD* | *M* | *SD* |
| HFRW | C3 | 4.38 | 1.65 | 0.72 | 0.23 | 0.77 | 0.22 | 0.83 | 0.25 | 0.74 | 0.24 |
|  | C4 | 4.05 | 1.66 | 0.82 | 0.37 | 0.88 | 0.41 | 0.88 | 0.32 | 0.77 | 0.29 |
| LFRW | C3 | 4.19 | 1.83 | 0.74 | 0.23 | 0.86 | 0.34 | 0.89 | 0.30 | 0.79 | 0.27 |
|  | C4 | 3.68 | 1.53 | 0.84 | 0.30 | 0.90 | 0.36 | 0.94 | 0.31 | 0.76 | 0.29 |
| HFPW | C3 | 4.18 | 1.61 | 0.79 | 0.20 | 0.80 | 0.31 | 0.82 | 0.20 | 0.87 | 0.32 |
|  | C4 | 4.10 | 1.66 | 0.76 | 0.27 | 0.78 | 0.24 | 0.84 | 0.26 | 0.83 | 0.27 |
| LFPW | C3 | 4.32 | 1.59 | 0.75 | 0.25 | 0.80 | 0.25 | 0.73 | 0.25 | 0.79 | 0.27 |
|  | C4 | 4.20 | 1.64 | 0.74 | 0.28 | 0.76 | 0.23 | 0.66 | 0.20 | 0.79 | 0.25 |

**Figure 2.**
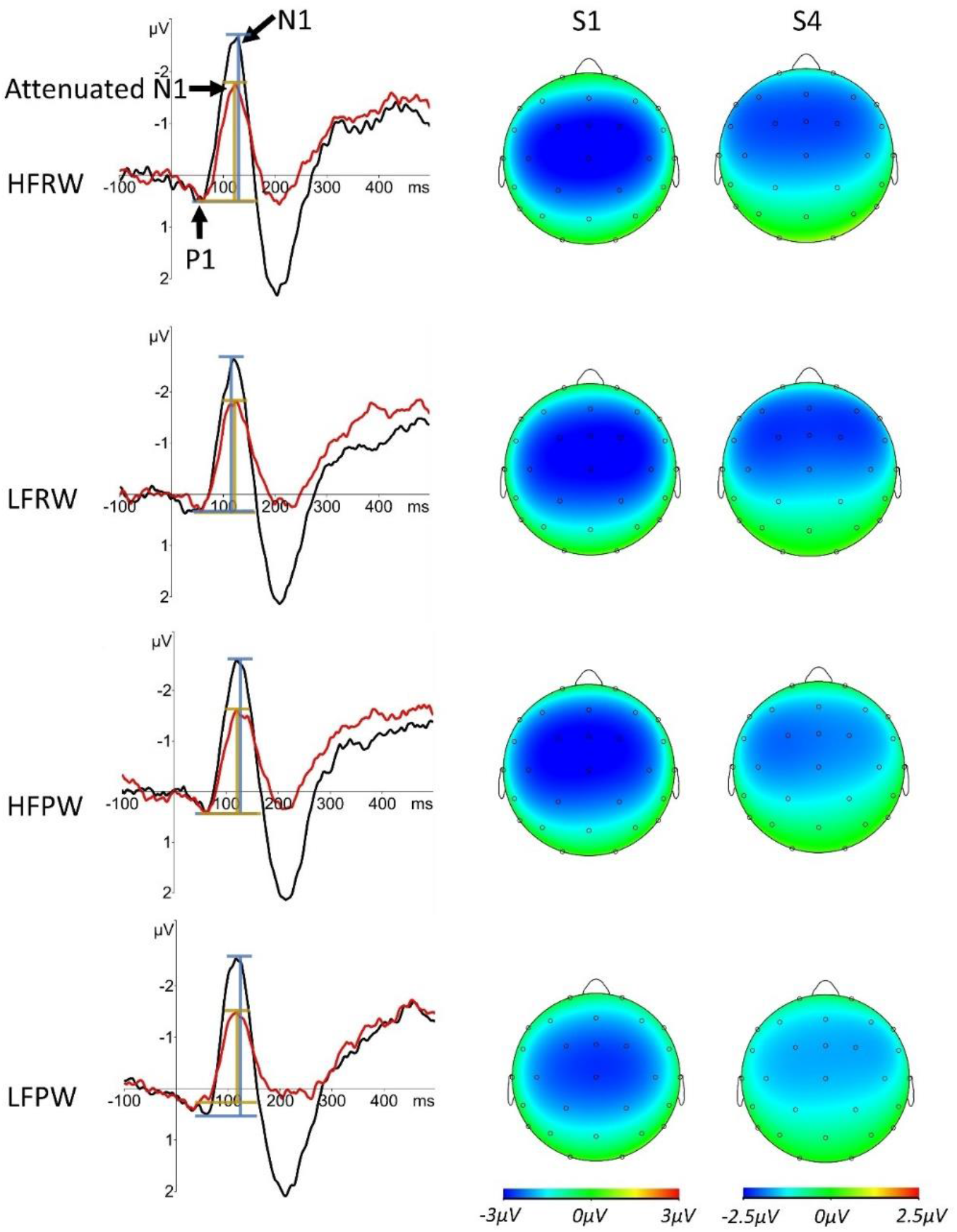
The grand average of the averaged ERP waveforms of C3 and C4 (left column) for the four conditions in S1 (black) and S4 (red), aligned with the topographic maps of the grand-averaged N1 according to the peak latency at C4 in S1 (middle column) and S4 (right column), in the four conditions. Differential scales were used for S1 and S4 for a demonstrative purpose. In the ERP waves, the blue bars denote the N1 peak-to-peak amplitudes (P1-N1) in S1, compared with the attenuated N1, as marked by the golden bars. Note the relatively lower ratio of the length of the line segment for S4 in the length of the segment for S1 in low-frequency pseudo word-forms (LFPW) than in high-frequency pseudo word-forms (HFPW) and in low-frequency real word-forms (LFRW), respectively.

**Figure 3.**
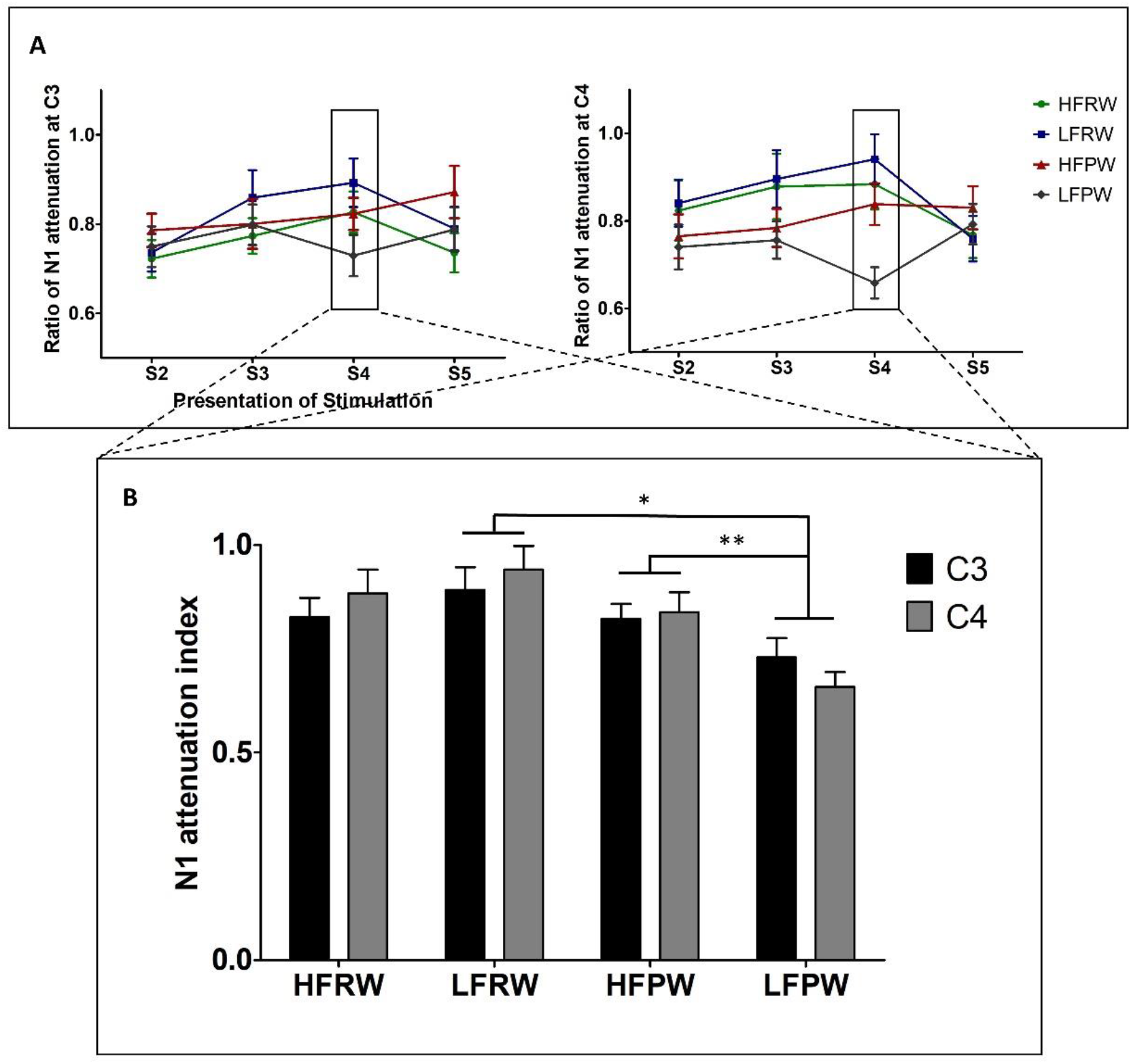
The phonological modulation effects of repetition-attenuation of N1. Panel A demonstrates the N1 repetition-attenuation indexes in the four conditions, in each stimulation position, at C3 and C4. Note the distinctive patterns of attenuation trajectories between real and pseudo word-forms. Panel B depicts the N1 repetition-attenuation indexes in S4 and the significant interactive effects between lexicality and frequency. The error bars represent the standard error of the mean (SEM), ** *p* < .01, *** *p* < .001

The inferential statistical analyses with the attenuation index data first revealed a main effect of LEXCIALITY (*F*_(1, 29)_ = 9.249, *p* = 0.02, partial *η*^2^ = 0.242) and a significant interaction between LEXICALITY and FREQUENCY (*F*_(1, 29)_ = 10.018, *p* = 0.016, partial *η*^2^ = 0.257) only in S4, after Bonferroni correction (Figure 2). Following this interaction, a main effect of LEXICALITY was found for low-frequency real and pseudo word-forms (*F*_(1, 29)_ = 16.854, *p* < 0.001, partial *η*^2^ = 0.368), as well as an interaction between LEXICALITY and ELECTRODE (*F*_(1, 29)_ = 5.419, *p* = 0.027, partial *η*^2^ = 0.157). Further analyses revealed main effects of LEXICALITY at both C3 (*F*_(1, 29)_ = 6.762, *p* = 0.015, partial *η*^2^ = 0.189) and C4 (*F*_(1, 29)_ = 24.432, *p* < 0.0001, partial *η*^2^ = 0.457). These results attest that the degree of N1 attenuation in low-frequency pseudo word-forms around the vertex is greater relative to that in low-frequency real word-forms. The effect size in C3 is about 16% and that in C4 is 28% (C3: LFPW *M* = 0.73, *SD* = 0.25, LFRW *M* = 0.89, *SD* = 0.30; C4: LFPW *M* = 0.66, *SD* = 0.20, LFRW *M* = 0.94, *SD* = 0.31) (See Table 1 and Figure 2).

Furthermore, unpacking the interaction between LEXICALITY and FREQUENCY also yielded a main effect of FREQUENCY for the two conditions of pseudo word-forms (*F*_(1, 29)_ = 9.793, *p* = 0.004, partial *η*^2^ = 0.252). This main effect confirms that the degree of N1 repetition-attenuation in low-frequency pseudo word-forms is about 9% greater than in high-frequency pseudo word-forms at C3 (LFPW: *M* = 0.73, *SD* = 0.25; HFPW: *M* = 0.82, *SD* = 0.20) and 18% greater at C4 (LFPW: *M* = 0.66, *SD* = 0.20; HFPW: *M* = 0.84, *SD* = 0.26) (Figure 2 & 3).

*Post hoc* analyses were conducted to test for the possibility that the phonological effects of N1 attenuation were decided by the P1, N1, and/or N1 (i.e., P1-N1) responses alone in S1 or S4, instead of the changes through stimulus repetitions. To this end, repeated measures ANOVA was conducted on the peak amplitudes of P1 and N1 component, and the peak-to-peak amplitudes of N1/P1 with three factors: LEXICALITY, FREQUENCY, and ELECTRODE, at S1 and S4, respectively. It is because only by comparing the N1 attenuation involving these two positions were phonological modulation effects identified. Neither main effects of LEXICALITY nor interactions between LEXICALITY and FREQUENCY were found in the analysis with the peak amplitudes at the two presentation positions. These results suggest that the neural decrement patterns reported in this study are not a reflection of the auditory processing of spoken words, but are decided by the attenuation modulated by differences in neural representations of phonological features.

## 4. Discussion

Although it is common to observe attenuated N1 responses of auditory ERPs to repetitively presented auditory stimuli, less is known about the role of cognitive/representational manipulations of auditory stimuli in influencing such physiological process. To address this issue, we tested the hypothesised modulation of the representations of two phonological features in spoken word-forms, namely lexicality and usage frequency, on N1 repetition-attenuation. The N1 data first attested that the current design could elicit decreased N1 responses throughout repeated stimulations. Then, we found evidence that the two phonological features are indeed factors that modulate the N1 repetition-attenuation, supporting our representational modulation hypothesis. To be specific, lexicality and frequency are interactive modulators on N1 repetition-attenuation. We found greater N1 repetition-attenuation in low-frequency pseudo word-forms than in low-frequency real word-forms (lexicality effect), and in high-frequency pseudo word-forms (frequency effect), respectively, at both C3 and C4 electrodes. These effects were identified at the S4 position in the stimulation trains, which may be attributed to the special recovery cycles of repetition attenuation for the present auditory stimuli (*cf*. Budd et al., 1992; Sambeth et al., 2004; Wang & Knösche, 2013).

Additionally, the results of *post hoc* analysis ensure that none of these phonological effects are just a reflection of the processing effects as indicated by the peak amplitudes of N1 or its preceding positive component P1 at S1 or S4. Therefore, to the best of our knowledge, our finding in the current study is the first showing that the repetition-attenuation of auditory N1 is interactively modulated by different types of phonological representations of speech sounds.

### 4.1. The lexicality and frequency effects, and their interactions

Here, the lexicality effect of N1 repetition-attenuation is partly in line with a previous study (Yue et al., 2017). In that study, researchers found smaller decrement of auditory N1 through the repetitions of a real word relative to a pseudoword in Mandarin over the electrodes F4, FC4, and C4, forming a region of interest, located in the right hemisphere. In the present study, we adopted a similar experimental design but with multiple real and pseudo word-forms, and identified greater degree of N1 repetition-attenuation in repeated pseudo word-forms than in real word-forms when the word-forms were of low usage frequency, at the fourth presentation position in the stimulation trains (i.e., S4). Moreover, this effect was found at the two electrodes of interest defined *a priori*, namely C3 and C4. Although the lexicality effect in the current result seems to exhibit a broader scalp distribution than Yue et al. (2017) (i.e., not only at a recording site in the right scalp, but also in the left), it may not be necessary to treat them as differential N1 repetition-attenuation effects. The main reason is that, the brain is a volume conductor (Luck, 2005), and therefore, it cannot be certain that the electoral responses recorded at C3 and C4 over the scalp are produced by different generators. Consequently, the seemingly extended lexicality effect in the current study and the right scalp effect in the previous study may have similar originality which responds to repeated auditory stimulations. Besides the lexicality and frequency effects at C4 identified in the current study demonstrate greater effect sizes and higher reliability than those at C3, which echo with the right-hemispheric effect of lexicality reported in Yue et al. (2017).

Regarding the difference between the two lexicality effects, a tentative explanation may be the relatively lower presentation probability of target stimuli in the current study than in Yue et al. (2017). In that study, the presentation probability of the stimuli in a condition was about 78.6% (55 experimental trains plus 15 filler trains) in contrast with 17% in the current study (e.g., in a block, 11 trains for each of the four conditions (44 trains in total) and 20 filler trains). As a result, the relatively lower presentation probability of each type of the word-forms in the current study might have induced differential neural adaptation than the higher presentation probability of the word-forms in Yue et al. (2017). This explanation is in line with some auditory discrimination studies, finding that the probability of deviant stimuli can either influence the MMN responses recorded from the scalp (e.g., Sabri & Campbell, 2001) or the electrophysiology recorded from single cell (e.g., Ayala & Malmierca, 2012). However, it must be pointed out that even if such differential adaptation matters for the scalp distribution of the lexicality effect, it does not hinder the sensitivity of the N1 repetition-attenuation to the lexical status of speech stimuli. This means that the attenuation of the N1 responses is indeed modulated by the phonological factor.

Moreover, the lexicality effect was only identified between real and pseudo word-forms with low frequency, suggesting not only involvement of usage frequency in modulating the N1 repetition-attenuation, but also a clear interplay between lexicality and frequency features. These results, together, are coherent with previous neurophysiological studies finding pre-attentive processing of phonological features such as lexicality (Pulvermüller, et al. 2001, Jacobsen et al., 2004) and lexical frequency (Aleksandrov et al., 2016), and, additionally, their interactions (Shtyrov et al., 2011; Silva et al., 2020). It can be suggested that the human brain can process multiple types of phonological features during pre-attentive speech perception, which eventually fuel the modulation on the repetition-attenuation of N1 in an interactive way.

In addition, the phonological effects are not likely to be explained by phonetic confounds — the tonal contrasts between real and pseudo word-forms with the same segmental templates, or the segmental differences between high and low frequency word-forms. Admittedly, there are studies reporting neurophysiological responses related to the pre-attentive processing of tonal and segmental features in Chinese language speakers (e.g., Luo et al., 2004; Malins & Joanisse, 2012; Wang et al., 2012). However, such accounts are unlikely to be applicable for the current results because of the reasons as follows. First, the phonological effects were hypothesised *a priori* based on previous studies. Particularly, the lexicality effect turns out to be a general replicate of a previous study (Yue et al., 2017) in which the effect was obtained by using a real (/ma1/) and a pseudo word-form (/na1/) that carried the same tone but different initial onset consonants. The similarity between the two lexicality effects obtained with materials that either containing tonal or segmental contrasts suggests that the smaller degree of N1 repetition-attenuation in real word-forms relative to pseudo word-forms can be caused by their representational difference, instead of the phonetic contrast.

Second, it is not sufficiently grounded on empirical findings to associate the phonological effects with tonal or segmental contrasts, in that few data have shown that repetition-attenuation of auditory N1 is sensitive to phonetic-level information. For example, also in Yue et al. (2017), in the two experimental (/ma1/ vs */na1/) and two control conditions (/mi2/ vs /ni2/), no effects related to tonal (Tone 1 vs Tone 2) or segmental features (/m/ vs. /n/) were identified.

Third, the design of the experiment made the phonetic account less likely to be suitable for the current data. Although the distributions of tones and segments in different conditions were not meant to be balanced, they were generally comparable. Therefore, we believe our design allowed us to test our hypothesis based on phonological representations before there is convincing evidence of phonetic modulation of auditory N1 to speech sounds. Nevertheless, further investigations on the phonological effects can be conducted by controlling the tonal or the segmental factors.

### 4.2. Theoretical implications

Our results provide insights into the underlying mechanism of the phonological/representational modulation on N1 repetition-attenuation. First, N1 repetition-attenuation is not likely a totally bottom-up, refractory process. Otherwise, no systematic difference should be observed during perceiving spoken word-forms that are acoustically similar but phonologically distinctive (Marklund et al., 2020). However, we reiterate our argument that it is not meant to refute the existence of refractoriness which may be a basic component of neural attenuation induced by any rapidly repeated auditory stimuli (Wang & Knösche, 2013), but to emphasise the modulatory role of phonological representations.

Besides, the phonological modulation on N1 repetition-attenuation is unlikely to be caused by high cognitive functions such as expectation (Todorovic et al., 2011) and selective attention (Öhman & Lader, 1972). This is first because that the participants were passively delivered with stimuli in different conditions by means of trains consisting of the same number of stimulation positions, separated by a constant interstimulus interval. Therefore, even if the participants had some expectations after being familiar with the experimental settings, such expectation should be equally applied for the N1 repetition-attenuation in all conditions. Furthermore, the participants’ attention was successfully distracted by a task which is unrelated with hearing. Hence, we can assume that the participants were in a stable attentional status during the experiment.

By excluding the potential physiological and cognitive accounts, the most possible explanation for the reliable N1 repetition-attenuation effects of lexicality and usage frequency is that, as is hypothesised, during the pre-attentive speech perception, the representational properties of phonological features are processed and modulate the repetition-attenuation of N1. This finding is, at least in part, coherent with the previous experiment finding that lexicality is a modulator of N1 short-term habituation (Yue et al., 2017). Since the representational difference between real and pseudo word-forms lies in the existence of lexical-level representations for real word-forms but only sublexical-level representations for pseudo word-forms, the current result indicates that N1 repetition-attenuation can capture the difference of representational levels of spoken word-forms. The N1-attenuation’s sensitivity to stimuli with contrastive representational levels has also been revealed by previous studies finding differential N1 repetition-attenuation patterns between acoustically matched speech and nonspeech sounds, that is, a higher representational level for speech sounds and a lower level for nonspeech sounds (Marklund et al., 2020; Teismann et al., 2004; Woods & Elmasian, 1986).

Second, apart from the lexicality modulation, the interaction between lexicality and frequency clearly suggests that the usage frequency is also a modulator of the N1 repetition-attenuation. Moreover, since the usage frequency in this study decides how word-forms are represented within the same representational level of phonology, this result suggests that representational modulations are not only sensitive to the between-levels features but also to within-level features. However, the within-level representational feature’s modulatory role must be played by interacting with the between-level feature. To be specific, in the lexical level (i.e., real word-forms), even though there is no difference between the N1 repetition-attenuation in high- and low-frequency real word-forms, the reliable lexicality effect is only observed in the comparison between low-frequency real- and pseudo-word-forms. This result clearly suggests that both of lexicality and frequency of low-frequency real word-forms were actually processed. In the sublexical level (i.e., pseudo word-forms), the consistent frequency effect in pseudo word-forms suggests clear classification between high- and low-frequency pseudo word-forms. Therefore, these findings together indicate that during perceiving repeated spoken-word input, the within-level representational feature leads to differential modulation outcomes, when its modulation is executed at different representational levels.

Such difference in the modulatory roles of within-level representation between the higher and lower representational levels of spoken word-forms may be ascribed to the top-down feedback from the lexical-level representations to the sublexical-level representations, given a widely accepted assumption of the hierarchical structure of phonological representations (Dahan & Magnuson, 2006). According to some modern models of speech perception such as TRACE (McClelland & Elman, 1986) or Predictive coding (Davis & Sohoglu, 2020), when a lexical entry exists in the lexicon, it has a corresponding lexical-level representation for the whole word. Then, the phonetic input can activate these representations to produce top-down feedback to the phonological processing at sublexical representational levels. In contrast, for pseudowords, there are no corresponding lexical entries and thus, the lack of top-down feedback results in differential neural and behavioural responses between pseudo and real word-forms. Such mechanism may be of particular interest of the current study as the interactive effect was found at S4 when the sensory system has been familiarised by three preceding repeated stimuli in a stimulation train. Unfortunately, our study does not allow us to further discuss whether the interplay between the within- and between-level representation in modulating the N1 repetition-attenuation is speech-specific or a general mechanism for the auditory/sensory system. To test this issue, future experiments should be conducted by involving both speech and nonspeech sounds (e.g., Marklund et al., 2021).

It should be noted that the phonological modulation is not equal to the processing of these features. To examine pre-attentive processes of phonological features, the neurophysiological indicators (e.g., MMN) are usually obtained by setting phonological contrasts between conditions (Shtyrov & Pulvermüller, 2007). In this case, neural effects could be associated with phonological processing. However, in the current study, the same stimulus was presented repeatedly in trains for fixed times at a constant ISI. Thus, no phonetic or phonological features need to be identified or even compared overtly, but the only cognitive processing is to map the word-form input onto the restored neural representations repetitively, and are supportive of our representational hypothesis of the modulatory role of phonological features.

Last but not the least, the lexicality and frequency effects in the current study may correspond to the brain function of sensory filtering during pre-attentive processing stage. Sensory filtering is considered as a brain function that filters out irrelevant information to protect the sensory system from being overloaded by repeated input (Font-Alaminos et al., 2020). Low-frequency pseudo word-forms are the least possible sound patterns of Mandarin speech, and therefore, very likely to be judged as “irrelevant” speech input, compared with the other two types of word-forms, namely low-frequency real word-forms and high-frequency pseudo word-form. Then it is reasonable to observe more suppressed N1 responses through the repetition of the ‘irrelevant’ speech input than other types of word-forms.

According to this reasoning, the current protocol exhibits potential to be developed into clinical tools to evaluate sensory function that filters repeated sounds. Sensory filtering has been found to be abnormal, as indexed by the evoked N1 responses or its attenuation, in patients with psychiatric (Kessler & Steinberg, 1989), neurological (Choi et al., 2016), and genetic diseases (Ethridge et al., 2016), as well as in children with autism spectrum disorder (Font-Alaminos et al., 2020). Therefore, the protocol of our study may be particularly useful for assessing the function of the auditory/sensory system which is related to speech perception. Furthermore, since no overt tasks were adopted, our protocol could be particularly suitable for populations who have difficulty in giving behavioural responses during tests, such as aging, young children, psychiatric, or neurological cohorts. In addition, the effects we identified here are only with two electrodes situated around the vertex (C3 and C4), making it more realistic to be operated in a clinical context. Admittedly, we don’t know whether such assessment of phonological relevance is executed according to the given experimental materials or just the previously developed phonological knowledge, which merits further examination.

## 5. Conclusion

The present study investigates whether and how lexicality and use frequency of spoken word-forms are two phonological features that modulate the repetition-attenuation of auditory N1. By using monosyllabic Mandarin word-forms, we found the first evidence that the two factors modulate the N1 repetition-attenuation in an interactive way. This finding suggests that the repetition-attenuation of N1 is sensitive to the representational properties of auditory stimuli, and may be a potentially useful tool to detect the brain function of speech information filtering in pre-attentive auditory perception.

## Author notes

This study is financially supported by the Grant for Junior Researchers from National Social Science Fund of China [16CYY024]. The first author thanks Prof. Ianthi Tsimpli for the inspiring discussions and her insights about this study. The first author also thanks the Fundamental Research Funds for the Central Universities and China Scholarship Council.

## Footnotes

1 Here, N1 is used as a common term to refer to the N1 component of both auditory evoked brain potential and event-related field. Despite of the fact that N1 has several sub-components, here it mainly refers to the N1b at the vertex.

2 An asterisk marked before a word-form designates pseudo word-form.

3 Mandarin tone is a lexical prosody which is superimposed over the entire sonorant portion of the onset and the rime. There are four typical tone in Mandarin, whose contours can be described as flat (Tone 1), rising (Tone 2), dipping (Tone 3), and falling (Tone 4).

## References

Ayala, Y. A., & Malmierca Dr., M. S. (2012). Stimulus-specific adaptation and deviance detection in the inferior colliculus. Frontiers in Neural Circuits, 6(NOV), 1–16. https://doi.org/10.3389/fncir.2012.00089

Aleksandrov, A. A., Memetova, K. S., Stankevich, L. N., Knyazeva, V. M., & Shtyrov, Y. (2020). Referent’s Lexical Frequency Predicts Mismatch Negativity Responses to New Words Following Semantic Training. Journal of psycholinguistic research, 49(2), 187–198. https://doi.org/10.1007/s10936-019-09678-3

Balota, D. A., Cortese, M. J., Sergent-Marshall, S. D., Spieler, D. H., & Yap, M. (2004). Visual word recognition of single-syllable words. Journal of experimental psychology. General, 133(2), 283–316. https://doi.org/10.1037/0096-3445.133.2.283

Boersma, P., & Weenink, D. (2013). Praat: Doing phonetics by computer (Version 5.3.39). Institute of Phonetic Sciences of the University of Amsterdam. Retrieved from http://www.praat.org

Bonte, M. L., Mitterer, H., Zellagui, N., Poelmans, H., & Blomert, L. (2005). Auditory cortical tuning to statistical regularities in phonology. Clinical Neurophysiology, 116(12), 2765–2774. https://doi.org/10.1016/j.clinph.2005.08.012

Boutros, N. N., Gjini, K., Urbach, H., & Pflieger, M. E. (2011). Mapping repetition suppression of the N100 evoked response to the human cerebral cortex. Biological Psychiatry, 69(9), 883–889. https://doi.org/10.1016/j.biopsych.2010.12.011

Budd, T. W., Barry, R. J., Gordon, E., Rennie, C., & Michie, P. T. (1998). Decrement of the N1 auditory event-related potential with stimulus repetition: Habituation vs. refractoriness. International Journal of Psychophysiology, 31(1), 51–68. https://doi.org/10.1016/S0167-8760(98)00040-3

Choi, W., Lim, M., Kim, J. S., & Chung, C. K. (2016). Habituation deficit of auditory N100m in patients with fibromyalgia. European Journal of Pain, 20(10), 1634–1643. https://doi.org/10.1002/ejp.883

Davis, M. H. & Sohoglu E. (2020) Three Functions of Prediction Error for Bayesian Inference in Speech Perception. In: Gazzaniga, M., Mangun R., & Poeppel D. (Eds), The Cognitive Neurosciences, 6th Edition. MIT Press, Camb, MA, USA.

Duanmu, S. (2002). The phonology of standard Chinese. Oxford, England: Oxford University Press.

Dufour, S. (2008). Phonological priming in auditory word recognition: when both controlled and automatic processes are responsible for the effects. Canadian Journal of Experimental Psychology, 62(1), 33–41. https://doi.org/10.1037/1196-1961.62.1.33

Ethridge, L. E., White, S. P., Mosconi, M. W., Wang, J., Byerly, M. J., & Sweeney, J. A. (2016). Reduced habituation of auditory evoked potentials indicate cortical hyper-excitability in fragile X syndrome. Translational Psychiatry, 6(4), e787–8. https://doi.org/10.1038/tp.2016.48

Font-Alaminos, M., Cornella, M., Costa-Faidella, J., Hervás, A., Leung, S., Rueda, I., & Escera, C. (2020). Increased subcortical neural responses to repeating auditory stimulation in children with autism spectrum disorder. Biological Psychology, 149(May 2019), 107807. https://doi.org/10.1016/j.biopsycho.2019.107807

Friston K. (2005). A theory of cortical responses. Philosophical transactions of the Royal Society of London. Series B, Biological sciences, 360(1456), 815–836. https://doi.org/10.1098/rstb.2005.1622

Fruhstorfer, H. (1971). Habituation and dishabituation of the human vertex response. Electroencephalography and Clinical Neurophysiology, 30(4), 306–312. https://doi.org/10.1016/0013-4694(71)90113-1

Fruhstorfer, H., Soveri, P., & Järvilehto, T. (1970). Short -term habituation of the auditory evoked response in man. Electroencephalography and Clinical Neurophysiology, 28(2), 153–161. https://doi.org/10.1016/0013-4694(70)90183-5

Gansonre, C., Højlund, A., Leminen, A., Bailey, C., & Shtyrov, Y. (2018). Task -free auditory EEG paradigm for probing multiple levels of speech processing in the brain. Psychophysiology, 55(11), 1–18. https://doi.org/10.1111/psyp.13216

Garrido, M. I., Kilner, J. M., Kiebel, S. J., Stephan, K. E., Baldeweg, T., & Friston, K. J. (2009). Repetition suppression and plasticity in the human brain. NeuroImage, 48(1), 269–279. https://doi.org/10.1016/j.neuroimage.2009.06.034

Glanzer, M., & Adams, J. K. (1985). The mirror effect in recognition memory. Memory & cognition, 13(1), 8–20. https://doi.org/10.3758/bf03198438

Grau, C., Fuentemilla, L., & Marco-Pallarés, J. (2007). Functional neural dynamics underlying auditory event-related N1 and N1 suppression response. NeuroImage, 36(3), 522–531. https://doi.org/10.1016/j.neuroimage.2007.03.027

Groves, P. M., & Thompson, R. F. (1970). Habituation: A dual-process theory. Psychological Review, 77(5), 419–450. https://doi.org/10.1037/h0029810

Gu, F., Li, J., Wang, X., Hou, Q., Huang, Y., & Chen, L. (2012). Memory traces for tonal language words revealed by auditory event-related potentials. Psychophysiology, 49(10), 1353–1360. 10.1111/j.1469-8986.2012.01447.x

Herrmann, B., Maess, B., & Johnsrude, I. S. (2018). Aging affects adaptation to sound-level statistics in human auditory cortex. Journal of Neuroscience, 38(8), 1989–1999. https://doi.org/10.1523/JNEUROSCI.1489-17.2018

Howard, D., Patterson, K., Wise, R., Brown, W. D., Friston, K., Weiller, C., & Frackowiak, R. (1992). The cortical localization of the lexicons: Positron emission tomography evidence. Brain, 115(6), 1769–1782. https://doi.org/10.1093/brain/115.6.1769

Jacobsen, T., Horváth, J., Schröger, E., Lattner, S., Widmann, A., & Winkler, I. (2004). Pre-attentive auditory processing of lexicality. Brain and Language, 88(1), 54–67. https://doi.org/10.1016/S0093-934X(03)00156-1

Joos, K., Gilles, A., Van de Heyning, P., De Ridder, D., & Vanneste, S. (2014). From sensation to percept: The neural signature of auditory event-related potentials. Neuroscience and Biobehavioral Reviews, 42, 148–156. https://doi.org/10.1016/j.neubiorev.2014.02.009

Kessler, C., & Steinberg, A. (1989). Evoked potential variation in schizophrenic subgroups. Biological Psychiatry, 26(4), 372–380. https://doi.org/10.1016/0006-3223(89)90053-x

Luck, S. J. (2005). An introduction to the event-related potential technique. Cambridge, MA: MIT Press.

Luo, H., Ni, J. T., Li, Z. H., Li, X. O., Zhang, D. R., Zeng, F. G., & Chen, L. (2006). Opposite patterns of hemisphere dominance for early auditory processing of lexical tones and consonants. Proceedings of the National Academy of Sciences of the United States of America, 103(51), 19558–19563. https://doi.org/10.1073/pnas.0607065104

Malins, J. G., & Joanisse, M. F. (2012). Setting the tone: An ERP investigation of the influences of phonological similarity on spoken word recognition in Mandarin Chinese. Neuropsychologia, 50(8), 2032–2043. https://doi.org/10.1016/j.neuropsychologia.2012.05.002

Marklund, E., Gustavsson, L., Kallioinen, P., & Schwarz, I. C. (2020). N1 Repetition-Attenuation for Acoustically Variable Speech and Spectrally Rotated Speech. Frontiers in Human Neuroscience, 14(October), 1–12. https://doi.org/10.3389/fnhum.2020.534804

Matsuzaki, N., Nagasawa, T., Juhász, C., Sood, S., & Asano, E. (2012). Independent predictors of neuronal adaptation in human primary visual cortex measured with high-gamma activity. NeuroImage, 59(2), 1639–1646. https://doi.org/10.1016/j.neuroimage.2011.09.014

McClaskey, C. M., Dias, J. W., Dubno, J. R., & Harris, K. C. (2018). Reliability of measures of N1 peak amplitude of the compound action potential in younger and older adults. Journal of Speech, Language, and Hearing Research, 61(9), 2422–2430. https://doi.org/10.1044/2018_JSLHR-H-18-0097

McClelland, J. L., & Elman, J. L. (1986). The TRACE model of speech perception. Cognitive Psychology, 18(1), 1–86. https://doi.org/10.1016/0010-0285(86)90015-0

McEnery, A. M., & Xiao, R. Z. (2004). The Lancaster Corpus of Mandarin Chinese: A corpus for monolingual and contrastive language study. In M. T. Lino, M. F. Xavier, F. Ferreira, R. Costa, R. Silva (Eds), Proceedings of the Fourth International Conference on Language Resources and Evaluation (LREC) 2004 (pp. 1175–1178). Lisbon, Portugal: Centro Cultural de Belem.

Mildner, V. (2008). The cognitive neuroscience of human communication. New York, NY: Lawrence Erlbaum Associates.

Muenssinger, J., Stingl, K. T., Matuz, T., Binder, G., Ehehalt, S., & Preissl, H. (2013). Auditory habituation to simple tones: Reduced evidence for habituation in children compared to adults. Frontiers in Human Neuroscience, 7(JUL), 1–7. https://doi.org/10.3389/fnhum.2013.00377

Myers, E. B., Blumstein, S. E., Walsh, E., & Eliassen, J. (2009). Inferior frontal regions underlie the perception of phonetic category invariance. Psychological Science, 20(7), 895–903. https://doi.org/10.1111/j.1467-9280.2009.02380.x

Näätänen, R. (1995). The mismatch negativity: A powerful tool for cognitive neuroscience. Ear and Hearing, 16, 6–18. doi:10.1097/00003446-199502000-00002

Näätänen, R., & Picton, T. (1987). The N1 wave of the human electric and magnetic response to sound: a review and an analysis of the component structure. Psychophysiology, 24(4), 375–425. https://doi.org/10.1111/j.1469-8986.1987.tb00311.x

Öhman, A., & Lader, M. (1972). Selective attention and “habituation” of the auditory averaged evoked response in humans. Physiology and Behavior, 8(1), 79–85. https://doi.org/10.1016/0031-9384(72)90132-1

Orfanidou, E., Marslen-Wilson, W. D., & Davis, M. H. (2006). Neural response suppression predicts repetition priming of spoken words and pseudowords. Journal of Cognitive Neuroscience, 18(8), 1237–1252. https://doi.org/10.1162/jocn.2006.18.8.1237

Perea, M., Rosa, E., & Gómez, C. (2005). The frequency effect for pseudowords in the lexical decision task. Perception and Psychophysics, 67(2), 301–314. https://doi.org/10.3758/BF03206493

Pulvermüller, F., Kujala, T., Shtyrov, Y., Simola, J., Tiitinen, H., Alku, P., Alho, K., Martinkauppi, S., Ilmoniemi, R. J., & Näätänen, R. (2001). Memory traces for words as revealed by the mismatch negativity. NeuroImage, 14(3), 607–616. https://doi.org/10.1006/nimg.2001.0864

Rankin, C. H., Abrams, T., Barry, R. J., Bhatnagar, S., Clayton, D. F., Colombo, J., … Thompson, R. F. (2009). Habituation revisited: An updated and revised description of the behavioral characteristics of habituation. Neurobiology of Learning and Memory, 92, 135–138. doi:10.1016/j.nlm.2008.09.012

Reder, L. M., Nhouyvanisvong, A., Schunn, C. D., Ayers, M. S., Hiraki, K., Angstadt, P., Schools, P. P., Hiraki, K., Society, P., Society, S., Reder, L. M., & Mellon, C. (2000). A Mechanistic Account of theMirror Effect for Word Frequency : A Computational of Remember-ow Judgments in a Continuous Recognition Paradigm. 26(2), 294–320. https://doi.org/10.1037//0278-7393.26.2.294

Ritter, W., Vaughan, H. G., & Costa, L. D. (1968). Orienting and habituation to auditory stimuli: A study of short terms changes in average evoked responses. Electroencephalography and Clinical Neurophysiology, 25(6), 550–556. https://doi.org/10.1016/0013-4694(68)90234-4

Rosburg, T., Haueisen, J., & Kreitschmann-Andermahr, I. (2004). The dipole location shift within the auditory evoked neuromagnetic field components N100m and mismatch negativity (MMNm). Clinical Neurophysiology, 115(4), 906–913. https://doi.org/10.1016/j.clinph.2003.11.039

Rosburg, T., Haueisen, J., & Sauer, H. (2002). Habituation of the auditory evoked field component N100m and its dependence on stimulus duration. Clinical Neurophysiology, 113(3), 421–428. https://doi.org/10.1016/S1388-2457(01)00727-1

Rosburg, T., Zimmerer, K., & Huonker, R. (2010). Short-term habituation of auditory evoked potential and neuromagnetic field components in dependence of the interstimulus interval. Experimental Brain Research, 205(4), 559–570. https://doi.org/10.1007/s00221-010-2391-3

Sabri, M., & Campbell, K. B. (2001). Effects of sequential and temporal probability of deviant occurrence on mismatch negativity. Cognitive Brain Research, 12(1), 171–180. https://doi.org/10.1016/S0926-6410(01)00026-X

Sambeth, A., Maes, J. H. R., Quiroga, R. Q., & Coenen, A. M. L. (2004). Effects of stimulus repetitions on the event-related potential of humans and rats. International Journal of Psychophysiology, 53(3), 197–205. https://doi.org/10.1016/j.ijpsycho.2004.04.004

Sara, Y., Mozhayeva, M. G., Liu, X., & Kavalali, E. T. (2002). Fast vesicle recycling supports neurotransmission during sustained stimulation at hippocampal synapses. Journal of Neuroscience, 22(5), 1608–1617. https://doi.org/10.1523/jneurosci.22-05-01608.2002

Shiffrin, R. M., & Steyvers, M. (1997). A model for recognition memory: REM-retrieving effectively from memory. Psychonomic bulletin & review, 4(2), 145–166. https://doi.org/10.3758/BF03209391

Shtyrov, Y., & Lenzen, M. (2017). First-pass neocortical processing of spoken language takes only 30 msec: Electrophysiological evidence. Cognitive Neuroscience, 8(1), 24–38. https://doi.org/10.1080/17588928.2016.1156663

Shtyrov, Y., & Pulvermüller, F. (2007). Language in the Mismatch Negativity Design. Journal of Psychophysiology, 21, 176–187.

Shtyrov, Y., Kimppa, L., Pulvermüller, F., & Kujala, T. (2011). Event -related potentials reflecting the frequency of unattended spoken words: A neuronal index of connection strength in lexical memory circuits? NeuroImage, 55(2), 658–668. https://doi.org/10.1016/j.neuroimage.2010.12.002

Shu, H., Chen, X., Anderson, R. C., Wu, N., & Xuan, Y. (2003). Properties of school Chinese: implications for learning to read. Child development, 74(1), 27–47. https://doi.org/10.1111/1467-8624.00519

Shuster, L. I. (2009). The effect of sublexical and lexical frequency on speech production: An fMRI investigation. Brain and Language, 111(1), 66–72. https://doi.org/10.1016/j.bandl.2009.06.003

Silva, S., Vigário, M., Fernandez, B. L., Jerónimo, R., Alter, K., & Frota, S. (2019). The sense of sounds: Brain responses to phonotactic frequency, phonological grammar and lexical meaning. Frontiers in Psychology, 10, 1–11. https://doi.org/10.3389/fpsyg.2019.00681

Slowiaczek, L. M., & Hamburger, M. (1992). Prelexical facilitation and lexical interference in auditory word recognition. Journal of Experimental Psychology: Learning, Memory, and Cognition,18, 1239–1250. https://doi.org/10.1037/0278-7393.18.6.1239

Swink, S., & Stuart, A. (2012). Auditory long latency responses to tonal and speech stimuli. Journal of Speech, Language, and Hearing Research, 55(2), 447–459. https://doi.org/10.1044/1092-4388(2011/10-0364)

Teismann, I. K., Sörös, P., Manemann, E., Ross, B., Pantev, C., & Knecht, S. (2004). Responsiveness to repeated speech stimuli persists in left but not right auditory cortex. NeuroReport, 15(8), 1267–1270. https://doi.org/10.1097/01.wnr.0000129856.58404.2d

Todorovic, A., van Ede, F., Maris, E., & de Lange, F. P. (2011). Prior expectation mediates neural adaptation to repeated sounds in the auditory cortex: An MEG study. Journal of Neuroscience, 31(25), 9118–9123. https://doi.org/10.1523/JNEUROSCI.1425-11.2011

Toyokura, M. (2003). Influence of stimulus intensity on waveform of sympathetic skin response evoked by magnetic stimulation. Clinical Neurophysiology, 114(8), 1423–1430. https://doi.org/10.1016/s1388-2457(03)00162-7

Wang, X. D., Gu, F., He, K., Chen, L. H., & Chen, L. (2012). Preattentive extraction of abstract auditory rules in speech sound stream: A mismatch negativity study using lexical tones. PLoS ONE, 7(1), 1–7. https://doi.org/10.1371/journal.pone.0030027

Wiener, S., & Ito, K. (2015). Do syllable-specific tonal probabilities guide lexical access? Evidence from Mandarin, Shanghai and Cantonese speakers. Language, Cognition and Neuroscience, 30(9), 1048–1060. https://doi.org/10.1080/23273798.2014.946934

Wiener, S., & Turnbull, R. (2016). Constraints of Tones, Vowels and Consonants on Lexical Selection in Mandarin Chinese. Language and Speech, 59(1), 59–82. https://doi.org/10.1177/0023830915578000

Woods, D. L., & Elmasian, R. (1986). The habituation of event-related potentials to speech sounds and tones. Electroencephalography and Clinical Neurophysiology/Evoked Potentials, 65(6), 447–459. https://doi.org/10.1016/0168-5597(86)90024-9

Yue, J. (2016). Tone-word recognition in Mandarin Chinese: Influences of lexical-level representations. University of Groningen.

Yue, J., Alter, K., Howard, D., & Bastiaanse, R. (2017). Early access to lexical-level phonological representations of Mandarin word-forms: evidence from auditory N1 habituation. Language, Cognition and Neuroscience, 32(9), 1148–1163. https://doi.org/10.1080/23273798.2017.1290261

Zhang, Q., Zhang, J. X., & Kong, L. (2009). An ERP study on the time course of phonological and semantic activation in Chinese word recognition. International Journal of Psychophysiology, 73(3), 235–45. 10.1016/j.ijpsycho.2009.04.001

Ziegler, J. C., Tan, L. H., Perry, C., & Montant, M. (2000). Phonology Matters: The Phonological Frequency Effect in Written Chinese. Psychological Science, 11(3), 234–238. https://doi.org/10.1111/1467-9280.00247

